# A joint Bayesian framework for modeling *Plasmodium vivax* transmission

**DOI:** 10.64898/2026.04.07.717120

**Authors:** Legesse Alamerie Ejigu, Wakweya Chali, Teun Bousema, Chris Drakeley, Fitsum G. Tadesse, John Bradley, Jordache Ramjith

## Abstract

Plasmodium vivax transmission from humans to mosquitoes depends on the density of gametocytes that in turn depends on asexual parasite replication and gametocyte commitment. These processes are often analyzed separately, despite being biologically linked and measured with substantial uncertainty. We used a joint Bayesian latent-variable model to simultaneously analyze parasite density, gametocyte density, and mosquito infectivity while accounting for measurement error and propagating uncertainty across linked processes. The model was applied to individual-level data from three P. vivax transmission studies conducted in Ethiopia (n = 455). A tenfold increase in gametocyte density was associated with more than a twofold increase in the odds of mosquito infection (odds ratio [OR] = 2.32, 95% credible interval [CrI]: 2.12–2.54). Asexual parasite density was also positively associated with infectivity after accounting for gametocyte density (OR = 1.74, 95% CrI: 1.60–1.90), and inclusion of parasite density improved predictive performance. Pathway decomposition within the joint model indicated that approximately 41% of the parasite–infectivity association operated through gametocyte density. Increasing age was associated with lower asexual parasite density but higher gametocyte density, resulting in minimal overall association with infectivity. Predicted infection probability increased sigmoidally with gametocyte density, remaining low at lower densities before increasing sharply and approaching a plateau at higher densities. Gametocyte density produced the largest predicted changes in the proportion of infected mosquitoes, while asexual parasite density added predictive information not fully captured by measured gametocyte density alone. This approach links molecular parasite measurements with mosquito infection risk while accounting for measurement uncertainty and provides an interpretable framework for studying the P. vivax infectious reservoir.

**Author Summary:** Malaria transmission occurs when mosquitoes ingest sexual-stage parasites, called gametocytes, during a blood meal. In *Plasmodium vivax* infections, human-to-mosquito transmission depends on linked biological stages, including asexual parasite replication, gametocyte production, and mosquito infection. These processes are closely connected and often measured with uncertainty, making them difficult to study using standard approaches that analyze them separately. In this study, we applied a joint Bayesian model that analyzes parasite density, gametocyte density, and mosquito infectivity together while accounting for uncertainty in laboratory measurements. Using data from three studies in Ethiopia, we quantified how parasite density, gametocyte density, and host characteristics relate to mosquito infection. The analysis showed that measured gametocyte density alone did not fully explain variation in infectivity, and that asexual parasite density provided additional predictive information. We also found that age was associated differently with asexual parasite and gametocyte densities, resulting in little overall association with infectivity. This approach helps link molecular parasite measurements with transmission outcomes and improves understanding of the *P. vivax* infectious reservoir in endemic settings.

## Introduction

Malaria remains a major global health challenge, with an estimated 282 million cases and 610,000 deaths in 2024[1]. Among the *Plasmodium* species infecting humans, *Plasmodium vivax* is the most geographically widespread, contributing substantially to morbidity and posing unique challenges for elimination due to its ability to form dormant hypnozoites in the liver, which cause relapsing infections [2, 3]. Transmission from humans to mosquitoes is determined by the presence of transmissible gametocytes, which arise from preceding asexual parasite replication. In *Plasmodium vivax* infections, asexual parasite replication gives rise to gametocytes, which are subsequently ingested by mosquitoes during blood feeding and determine the probability of mosquito infection [3-6]. Quantifying how these within-host processes translate into transmission outcomes is essential for understanding the human infectious reservoir and for designing interventions aimed at reducing malaria transmission.

Accurate quantification of gametocytes, a key determinant of mosquito infectivity, remains a major challenge in P. vivax infections. Unlike *P. falciparum, P. vivax* gametocytes lack distinctive morphology [7] and often occur at low densities, making microscopy insensitive and prone to misclassification [8, 9]. Molecular assays, including transcript-based PCR methods, improve detection sensitivity and are increasingly used to relate gametocyte density to mosquito infection outcomes, but they still exhibit substantial variability[10]. These diagnostic limitations propagate uncertainty in estimates of transmission potential.

Quantifying *P. vivax* transmission is further complicated by the fact that infectiousness depends on multiple interdependent biological processes that are imperfectly observed. Human-to-mosquito transmission requires the presence of transmissible gametocytes, which arise from preceding asexual parasite replication. As a result, estimates of transmission potential often fail to reflect the underlying biological mechanisms that link within-host parasite dynamics to mosquito infection outcomes. This stage dependence, combined with typically lower parasite densities and diagnostic limitations, underscores the need for advanced modelling approaches to better characterize P. vivax transmission [11, 12].

Most statistical analyses of malaria transmission examine parasite density, gametocyte density, and mosquito infection outcomes as separate or conditionally independent processes. However, these quantities are biologically linked: gametocyte density arises from asexual parasite replication, and mosquito infection depends on the presence of transmissible gametocytes. In addition, parasite and gametocyte densities are often measured with substantial uncertainty due to sampling variability and limitations in molecular quantification methods. Analyzing these processes independently can therefore obscure their dependency structure and underestimate uncertainty in transmission estimates[6, 13]. In addition, while transmission is driven by gametocytes, asexual parasite density may still have residual associations with infectivity through host or vector responses unexplained by gametocyte densities, though such effects are difficult to estimate in standard regression frameworks. Asexual parasite density may influence mosquito infection indirectly by reflecting the parasite biomass from which gametocytes are produced, indicating infection duration, or improving estimation precision of gametocyte density. Conventional approaches also rarely account for measurement error, potentially leading to attenuation bias or overly precise estimates of infectivity [14].

Joint Bayesian latent-variable models provide a flexible way to simultaneously analyze interconnected biological processes that are imperfectly observed while propagating measurement uncertainty across stages of infection. Related joint models have been applied in other infectious disease settings, including linking viral load and immune responses in HIV [15, 16], antibody dynamics associated with risk in dengue [15], and combining time-to-infection with vector abundance to refine malaria transmission estimates[14]. Despite these advances in *P. vivax* transmission studies joint model remain underused where linked parasite stages and imperfect molecular measurements are central features of the data[3].

In this study, we used a joint Bayesian latent-variable model to simultaneously analyze asexual parasite density, gametocyte density, and mosquito infectivity in *P. vivax*. Using individual-level data from three malaria transmission studies conducted in Ethiopia, the model integrates measurement error and transmission components to quantify associations among parasite density, gametocyte density, and mosquito infection outcomes. This approach allows model parameters to be translated into predicted infection probabilities and differences in the proportion of infected mosquitoes under defined contrasts in host and parasite characteristics. Specifically, we aimed to (1) characterize the relationship between gametocyte density and mosquito infection probability, (2) quantify the extent to which the parasite–infectivity association operates through gametocyte density, and (3) evaluate how host factors, including age and fever status, relate to parasite density, gametocyte density, and transmission potential within a joint uncertainty-aware modeling framework.

## Methods

### Data

Data from three *P. vivax* transmission studies conducted in Adama and Arba Minch, Ethiopia were used. Study designs and participant selection procedures have been described elsewhere [17, 18]. Briefly, study one was a longitudinal cohort conducted in Arba Minch between 2020 and 2022; however, for consistency with other data sets, only baseline data from symptomatic individuals were included in the present analysis. Study two was a cross-sectional survey conducted in Adama between 2018 and 2019 among symptomatic self-presenting participants at health facilities. Study three, a cross-sectional study conducted in 2016 in Adama, included both symptomatic and asymptomatic individuals. All datasets contain individual-level parasitological and clinical measurements together with mosquito membrane feeding assays outcomes. A summary of the study characteristics, including collection years, sites, and sample sizes, is provided in the supplementary materials (S1 Table).

### Joint Bayesian model

A joint Bayesian latent-variable model that integrates asexual parasite density, gametocyte density, and mosquito infectivity was employed, explicitly accounting for measurement error and biological dependencies and implemented using the probabilistic programming language Stan[9, 19]. Observed log10 asexual parasite and gametocyte densities were treated as noisy measurements of underlying latent stage-specific densities. Latent asexual parasite density and latent gametocyte density were linked through a linear regression submodel reflecting their biological dependence. Mosquito infectivity outcomes were modelled using a beta-binomial likelihood with a logit link, allowing for overdispersion in mosquito infection outcomes. The full model comprises three components: (1) latent asexual parasite and gametocyte density, (2) measurement error sub models linking observed and latent quantities, and (3) a mosquito infection outcome model. This joint structure corrects for attenuation bias and propagates uncertainty across biological stages. Fig 1 summarizes the full model structure of the joint Bayesian latent-variable model, including the measurement layer linking observed molecular data to latent densities, the latent process layer capturing stage dependence, and the outcome layer relating latent states to mosquito infection outcomes.

**Fig 1.**
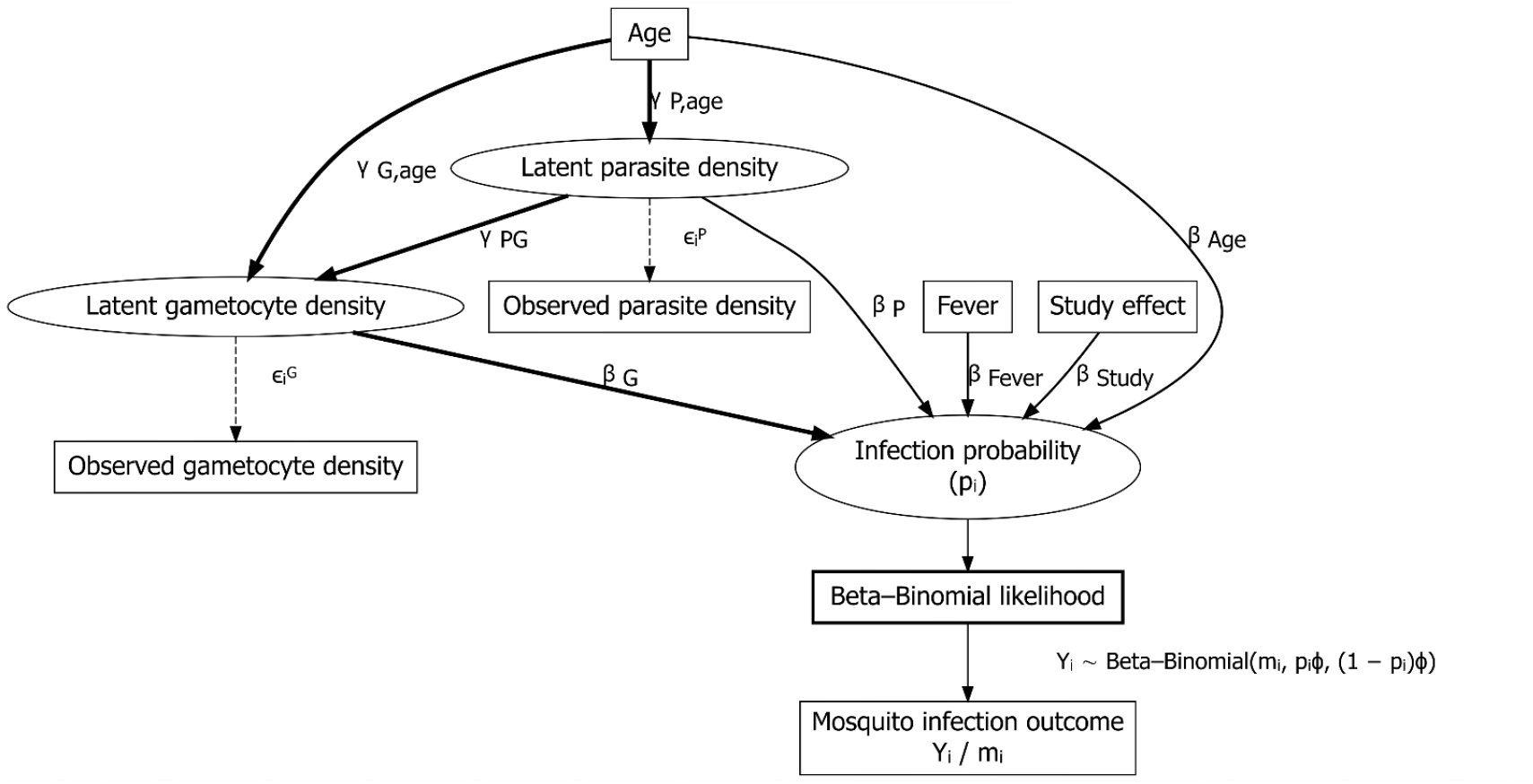
Graphical representation of the joint Bayesian latent-variable model. Observed asexual parasite density and gametocyte density are modeled as noisy measurements of the true (latent) parasite density 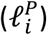 and gametocyte density 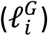, with measurement errors, respectively. Latent parasite density (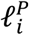 influences latent gametocyte density 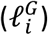) via the parameter( *γ*_*GP*_ ). Host age affects both latent parasite and gametocyte density, with effects denoted by *γ*_*Gage*_ and *γ*_*Page*_Latent gametocyte density, latent parasite density, age, fever, and study-level variability contribute to the infection probability (*p*_*i*_), which is linked to the observed mosquito infection outcome (*Y*_*i*_/*m*_*i*_) via a beta-binomial likelihood. Thick arrows represent key process links, dashed arrows represent measurement error relationships, and parameter symbols indicate estimated effects in the model.

### Model structure

#### Measurement error models

Let 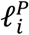 and 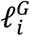 denote the true (latent) log-transformed asexual parasite and gametocyte densities for individual *i*, respectively. The observed values, 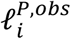 and 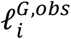, are then modeled as the true values plus random error:

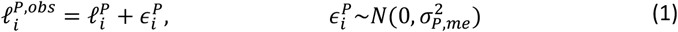

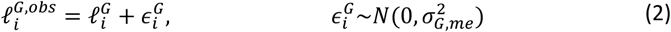

where, 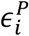 and 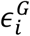 represent the measurement errors for parasite and gametocyte densities, respectively, and are assumed to follow a normal distribution with a mean of zero and variances 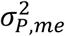 and 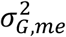. Errors are assumed to be independent within individuals. These variances quantify the degree of uncertainty associated with each measurement.

#### Latent process models

A linear regression framework was used to model the association between asexual parasite density and gametocyte density, as well as the associations of age with parasite and gametocyte density. This reflects the biological expectation that individuals with higher parasite burdens are more likely to have higher gametocyte densities, while recognizing that this relationship can be influenced by host and infection-related factors, including age and duration of infection, which is typically unobserved in cross-sectional measurements. Latent asexual parasite density is therefore modeled as a stochastic process to capture for residual between-individual biological variability beyond measured covariates:

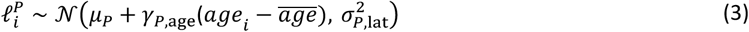

Latent gametocyte density is modeled conditional on latent parasite density:

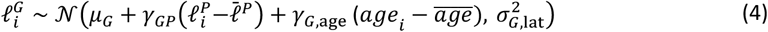

Where, *μ*_*p*_ and *μ*_*G*_ represent the (intercept) values for the actual parasite and gametocyte densities, when age is at its average value, respectively. 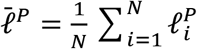 denotes the mean latent log asexual parasite density used for centering, *γ*_*GP*_ represents the association between parasite and gametocyte density, and *σ*_*P*,lat_ and *σ*_*G*,lat_ represent latent process variability.

#### Infection model

To model the association between latent gametocyte density and the proportion of infected mosquitoes while adjusting for asexual parasite density, fever, host age, and allowing study-specific variability, we used a beta-binomial likelihood to account for overdispersion.

Let *Y*_*i*_ be infected mosquitoes out of *m*_*i*_ dissected:

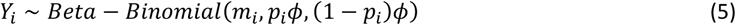

where *ϕ* is a precision parameter controlling overdispersion (larger values imply less variability around the binomial mean) the infection probability *p*_*i*_ is defined through a logit link:

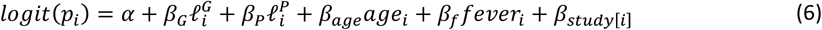

Here, *α* is the intercept, *β*_*G*_ *and β*_*P*_ represent latent gametocyte and asexual parasite density effects, *β*_*age*_ *and β*_*f*_ account for age and fever, and *β*_*study*[*i*]_ represents study-specific fixed effects, parameterized as indicator (dummy) variables, with one study treated as the reference category to ensure identifiability. Study effects were included as fixed intercept deviations in the infection model to adjust for systematic differences between studies, rather than modeled as random effects

#### Likelihood

The joint likelihood combines measurement and infection models. For the measurement models (eq. 1, 2), the likelihood for the observed density is:

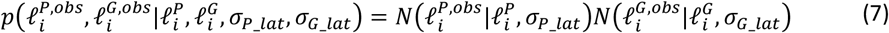

For the infection model **(5)**, the beta-binomial likelihood accounts for overdispersion in mosquito infection outcomes [20, 21]:

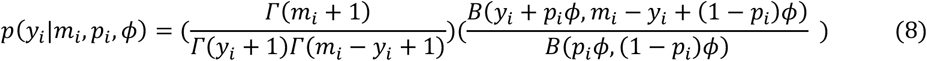

Where: *Γ* is the gamma function, B is the beta function, and *p*_*i*_ is defined by the logit model (eq. 6). This formulation captures the variability in *P. vivax* transmission beyond binomial assumptions [22].

The full likelihood is the product of all individuals:

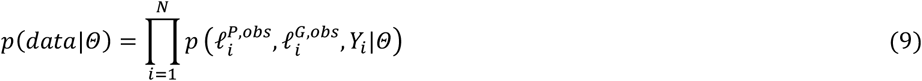

where *N* is the number of individuals and Θ denotes the full set of model parameters.

#### Prior distributions

Prior distributions were informed by *P. vivax* biology and Ethiopian studies [17, 18, 23], ensuring biological plausibility [24]. A detailed description of prior specifications and their justification is provided in Supplementary Table S2.

#### Bayesian inference and posterior estimation

Bayesian inference was performed using Markov Chain Monte Carlo (MCMC) sampling in Stan and fitted using CmdStanR [25]. The posterior distribution, proportional to the likelihood times the prior, was sampled to generate 8,000 draws (after burn-in), with medians and 95% credible intervals (2.5% and 97.5% quantiles) reported [20]. Convergence was confirmed using the Gelman-Rubin diagnostic 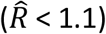 and effective sample sizes (>400 per chain). Posterior predictive checks generated replicated infection outcomes and log-likelihoods to assess model fit [21]. For interpretability, regression coefficients from the model were transformed to odds ratios.

#### Estimation of pathway-specific and total associations

To quantify how host factors (age, fever, and asexual parasite density) are associated with mosquito infectivity, we decomposed model-based associations into pathway-specific and residual conditional components. This decomposition follows the structure of pathway-based association analyses within a Bayesian framework[26, 27], but is interpreted here as describing conditional along biologically motivated pathways rather than causal mediation in the counterfactual sense. In the joint model, asexual parasite density and gametocyte density are represented as latent biological quantities measured with error, and mosquito infectivity is modeled as a downstream transmission outcome. The model structure reflects the biological ordering whereby asexual parasite density predicts gametocyte density, which in turn predicts infectivity. Within this framework, pathway-specific associations are defined algebraically as products of regression coefficients along the specified parasite-gametocyte-infectivity pathway. Residual conditional associations correspond to the regression coefficient of a covariate in the infectivity model after conditioning on the modeled parasite and gametocyte densities. Importantly, these quantities represent model-implied conditional associations rather than causal effects in the counterfactual sense. For asexual parasite density in particular, the direct effect should be interpreted as a residual conditional effect after accounting for gametocyte density. Since only gametocytes can infect mosquitoes biologically, this residual effect does not imply a separate transmission pathway, but instead likely captures unmeasured aspects of gametocyte biology (e.g., maturity, viability, or timing) that are correlated with asexual parasite density but not fully represented by measured gametocyte density.

The full decomposition of pathway-specific direct and indirect effects derived from the joint posterior distribution is described in the Supporting Information (Supplementary Methods). All quantities were computed for each MCMC posterior draw, and results were summarized using posterior means and 95% credible intervals (CrI). Given the cross-sectional nature of the data and the lack of temporal ordering, these decompositions should be interpreted as describing conditional associations along biologically motivated pathways rather than causal mediation in the counterfactual sense.

#### Prediction framework

To inform intervention strategies, we generated predictions across a grid of parasite and gametocyte density for all individuals, incorporating study-specific effects. Predicted infection probabilities, derived from posterior draws, provide uncertainty estimates for identifying ranges of parasite and gametocyte density associated with higher transmission risk[24].

#### Prediction and probability contrasts

To facilitate interpretation of model results, predicted mosquito infection probabilities were calculated under specific contrasts in model covariates. The effect size is summarized using the risk difference (RD).

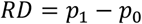

Where: *p*_0_ is the predicted probability of mosquito infection when all model covariates are fixed at specified values and p_1_ is the predicted probability after changing one covariate of interest while holding the remaining covariates fixed at the same values. Detailed descriptions of the methods are provided in S1 Text)

These contrasts quantify predicted changes in the proportion of infected mosquitoes under defined perturbations of parasite density, gametocyte density, age, and fever status. Predicted contrasts were calculated for each posterior draw and summarized by posterior medians and 95% credible intervals

## Results

A total of 455 individuals from three P. vivax transmission studies conducted in Ethiopia were included in the analysis. Of these, 304 participants (66.8%) were infectious to mosquitoes, with the highest proportion of infectious individuals was observed in Arba Minch (70.5% of participants infecting at least one mosquito) and the lowest in the Adama cohort (50.9%). The study population was predominantly male (67.5%). Age distributions varied by study site, with a higher proportion of participants under 15 years of age in Arba Minch than in Adama. Clinical presentation also differed by study design: over half of participants in Arba Minch reported fever, whereas fever was less common in Adama, particularly in the study that included asymptomatic individuals. On average, 30 mosquitoes per participant were dissected to assess infectivity, with minimal variation across sites (median: 30; IQR: 30-31) (Table 1).

**Table 1.**
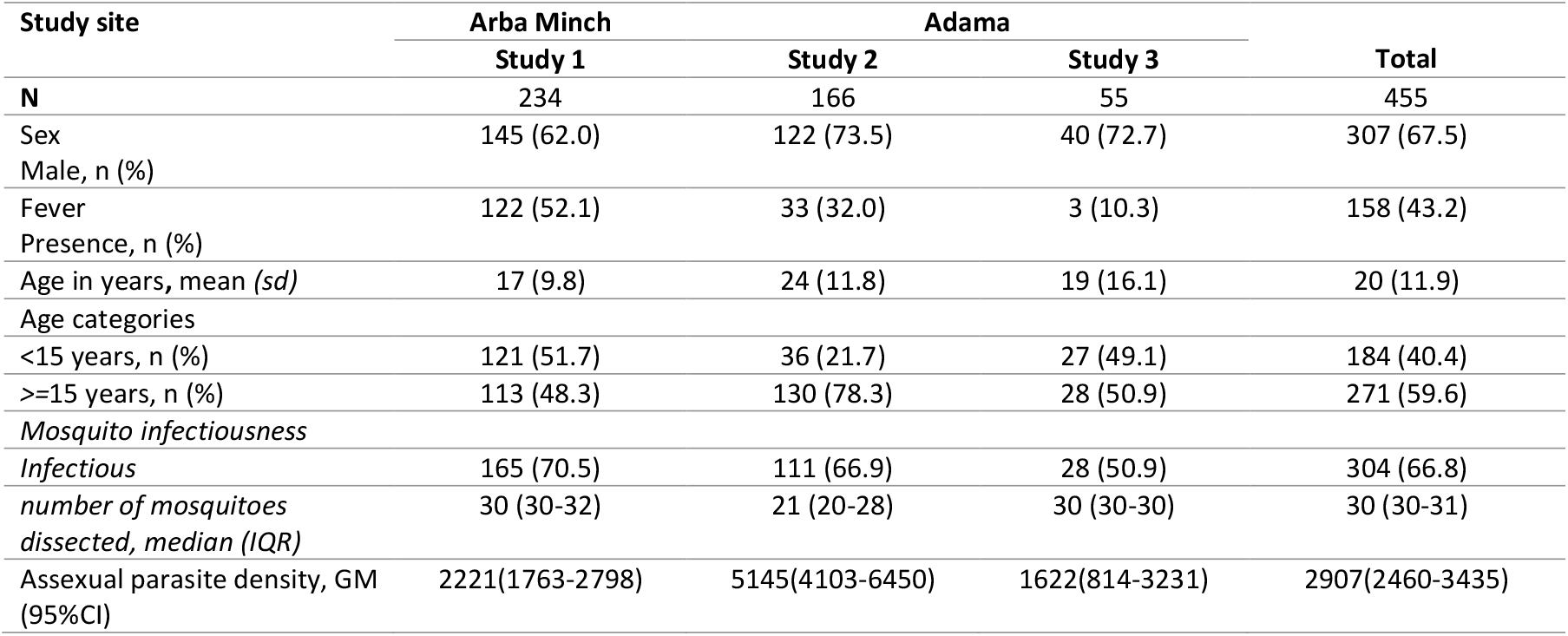
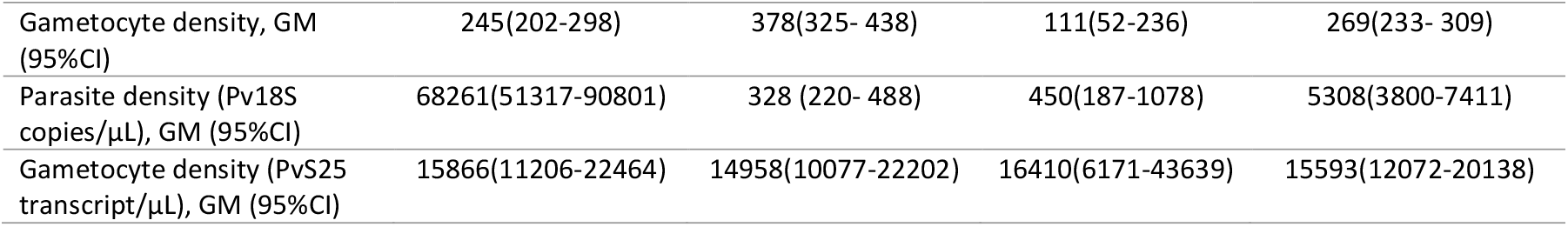
Demographic characteristics of participants by study and site.

### Parasite and gametocyte density distributions across study sites

Parasite density distributions differed significantly across the three studies (Kruskal-Wallis test, p < 0.001) (Fig 2A and 2B). Microscopy-based parasite and gametocyte counts were highest in Study 2 (Adama, 2018–2019), whereas qPCR-based *Pv18S* copies were notably elevated in Study 1 (Arba Minch, 2020–2022). Gametocyte density, quantified via *Pvs25* transcript levels, showed no significant difference between studies (p = 0.82). Substantial heterogeneity was observed at the individual level, consistent with the variable nature of *P. vivax* infections in field settings (Fig 2C and 2D).

**Fig 2.**
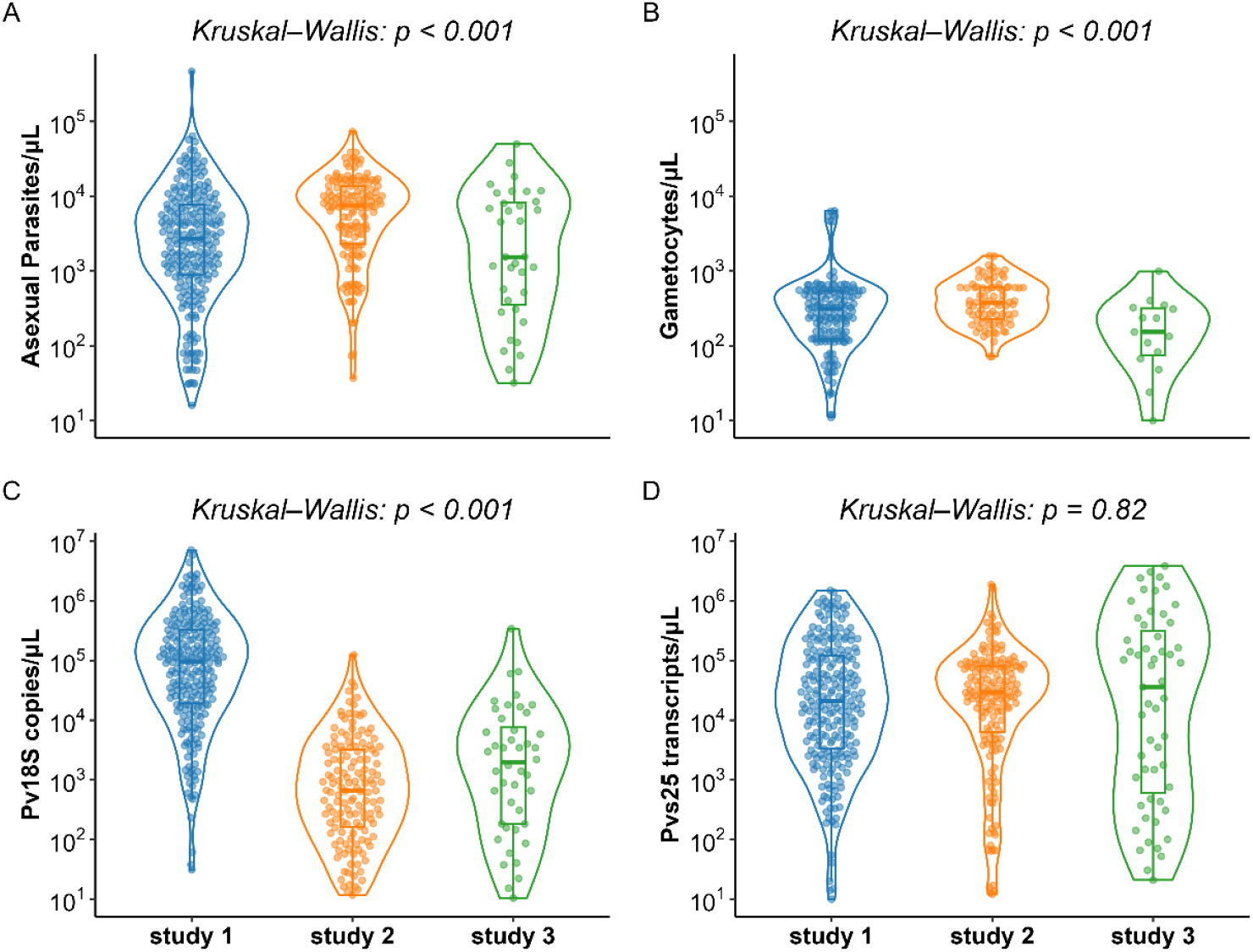
Distribution of parasite and gametocyte densities across three studies. Violin plots show the distributions of (A) asexual-stage parasite densities (parasites/μL), (B) gametocyte densities (gametocytes/μL), (C) Plasmodium vivax 18S rRNA gene copies/μL, and (D) P. vivax Pvs25 transcript copies/μL for each study dataset. Points represent individual observations overlaid on boxplots indicating the median and interquartile range. All values are displayed on a log10 scale. Colors indicate study: Study 1 (blue), Study 2 (orange), and Study 3 (light green).

### Correlation between parasite and gametocyte densities

A positive correlation was observed between parasite density (Pv18S copies/µL) and gametocyte density (Pvs25 transcripts/µL) across all studies (Fig 3). Spearman correlation coefficients ranged from moderate in Study 1 (ρ = 0.60, p < 0.001) to strong in Study 2 (ρ = 0.78, p < 0.001) and Study 3 (ρ = 0.85, p < 0.001). The positive association between parasite and gametocyte density was consistent across studies, although the strength of association varied. Age-stratified analysis (quintiles: Q1 = youngest 20%, Q3 = oldest 20%) showed that parasite density was highest in Q1 and declined progressively with age, whereas gametocyte density remained relatively stable across age groups. This pattern was visually apparent across all three study sites, with broadly similar age-related trends, although the magnitude of decline varied (Fig 3A, 3B, and 3C).

**Fig 3.**
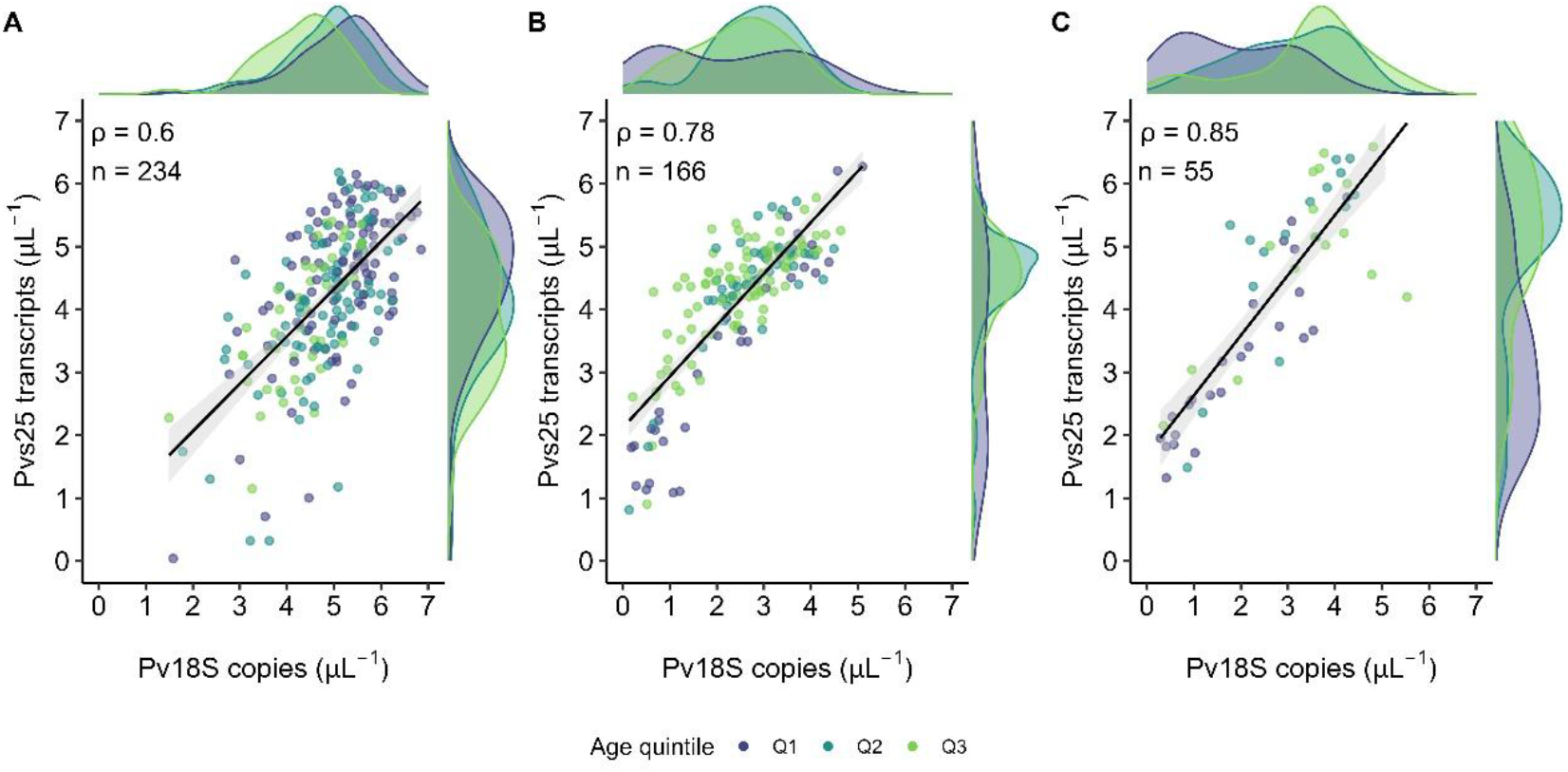
Association between P. vivax parasite (Pv18S copies/µL) and gametocyte densities (Pvs25 transcripts/µL), stratified by age quintiles (Q1-Q3). Points represent individual measurements, with colors indicating age groups (Q1: youngest 20%, Q3: oldest 20%). The solid line shows the linear regression fit with shading indicating the 95% confidence interval. The top and right figures indicate the density distribution by age quintiles by study (A=study 1, B=study 2, and C=study 3).

### Joint Bayesian model result

The joint Bayesian latent-variable model simultaneously estimated latent asexual parasite density, latent gametocyte density, and mosquito infection probability while accounting for measurement error and between-study heterogeneity. Latent densities refer to the underlying true parasite and gametocyte levels after accounting for measurement error. Model convergence diagnostics indicated good mixing and stable posterior estimates 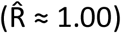, and posterior predictive checks showed good agreement between observed and predicted infection outcomes.

The conditional association between gametocyte density and infectivity was strong (β = 0.84, 95% CrI: 0.75–0.93), corresponding to an odds ratio (OR) of 2.32 (95% CrI: 2.12–2.54) for a tenfold increase in gametocyte density (Table 2 and Fig 4). Predicted infection probability increased sigmoidally with gametocyte density, with low predicted probability at lower gametocyte densities and a sharp increase across an estimated range where infection probability begins to increase around 100 transcripts/µL (95% CrI: 31–292) (Fig 5).

**Table 2.** Summary of parameter estimates from Joint Bayesian model. Estimates are posterior means with 95% credible intervals (CrI). ESS denotes effective sample size and 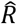 indicates convergence. Latent parasite and gametocyte densities represent underlying (error-corrected) biological quantities. Direct effects (*γ, β*) describe associations with parasite density, gametocyte density, and infectivity (log-odds scale). Mediation estimates include indirect effects (IE), representing associations operating through intermediate pathways (i.e., parasite → gametocyte → infectivity), and total effects (TE), representing overall associations across pathways.

| Parameter | Mean (95% CrI) | ESS (bulk) | $\hat{R}$ |
| --- | --- | --- | --- |
| **Intercepts** |  |  |  |
| $\alpha$ | -6.99 (-7.56, -6.43) | 30,598 | 1.00 |
| $\mu_P$ | 3.71 (3.57, 3.85) | 6,527 | 1.00 |
| $\mu_G$ | 4.17 (4.10, 4.25) | 18,532 | 1.00 |
| **Dispersion** |  |  |  |
| $\phi$ | 1.54 (1.35, 1.75) | 64,639 | 1.00 |
| **Latent standard deviation** |  |  |  |
| $\sigma_{P,lat}$ | 1.44 (1.34, 1.55) | 10,144 | 1.00 |
| $\sigma_{G,lat}$ | 0.85 (0.78, 0.93) | 13,662 | |
| **Direct effects on parasite and gametocyte densities** |  |  |  |
| $\gamma_{P\_age}$ | -0.03 (-0.04, -0.02) | 7,774 | 1.00 |
| $\gamma_{G\_age}$ | 0.02 (0.01, 0.03) | 19,611 | 1.00 |
| $\gamma_{GP}$ | 0.45 (0.39, 0.52) | 13,925 | 1.00 |
| **Infectivity estimates** |  |  |  |
| $\beta_G$ | 0.84 (0.75, 0.93) | 55,929 | 1.00 |
| $\beta_P$ | 0.56 (0.47, 0.64) | 40,273 | 1.00 |
| $\beta_f$ | -0.45 (-0.75, -0.15) | 63,220 | 1.00 |
| $\beta_{age}$ | 0.02 (0.01, 0.03) | 67,211 | 1.00 |
| **Mediation estimates** |  |  |  |
| IE (age → gametocyte) | -0.012 (-0.017, -0.0076) | 8,327 | 1.00 |
| TE (age → gametocyte) | 0.006 (-0.002, 0.013) | 13,827 | 1.00 |
| IE (age → infectivity) | -0.010 (-0.021, 0.0004) | 9,582 | 1.00 |
| TE (age → infectivity) | 0.008 (-0.007, 0.02) | 16,477 | 1.00 |
| IE (parasite → infectivity) | 0.38 (0.32, 0.44) | 18,567 | 1.00 |
| TE (parasite → infectivity) | 0.94 (0.85, 1.02) | 25,324 | 1.00 |

**Figure 4.**
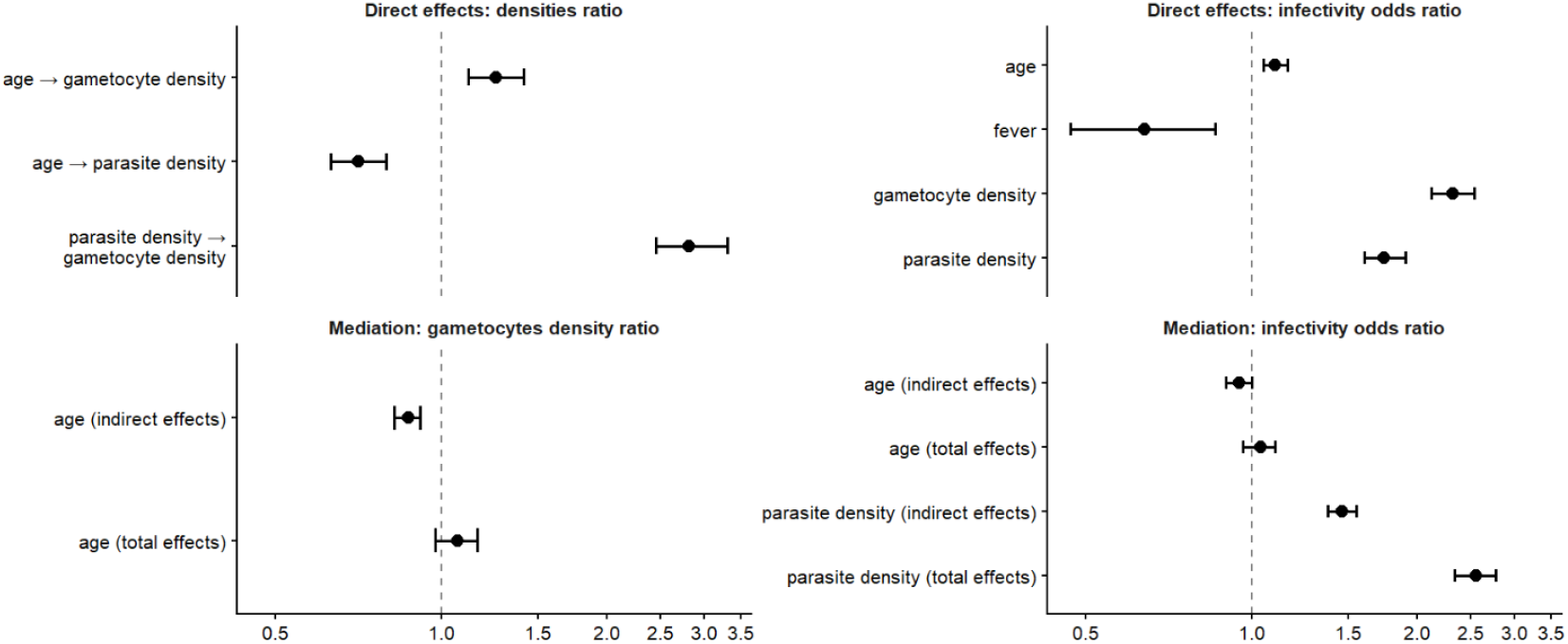
Forest plot of direct and mediated effects from the joint Bayesian transmission model. Points represent posterior means and horizontal bars indicate 95% credible intervals (CrI). Effects are shown as density ratios (for parasite and gametocyte sub models) and odds ratios (for infectivity sub model). The vertical dashed line at 1 denotes no association. Age effects are expressed by a 5-year increase. Mediation effects represent indirect pathways through gametocyte density, while direct effects represent associations after accounting for parasite and gametocyte densities. Density ratios are interpreted on the log10 scale (i.e. per tenfold increase), while odds ratios represent multiplicative changes in the odds of mosquito infection.

**Fig 5.**
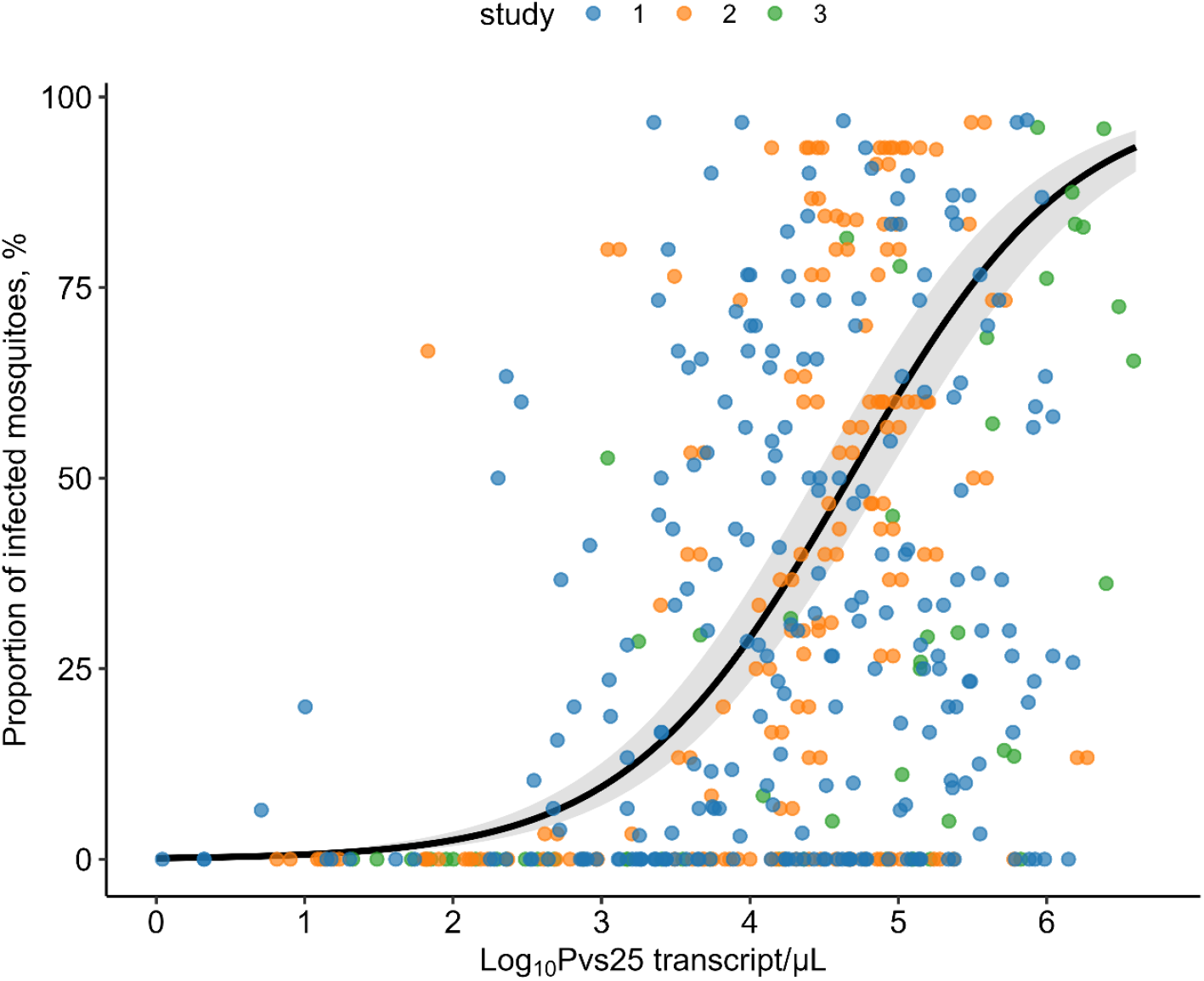
Predicted mosquito infection probability as a function of latent log10 gametocyte density. This figure presents the predicted relationship between the predicted mosquito infection probability and the latent log10 gametocyte density (logG), as estimated by a Bayesian joint error model. Point color denotes the observed proportion of mosquito infection from three different studies (blue = Study1, orange = Study2, and green = Study3). The black line displays the fitted relationship, and the shaded area around the line indicates the 95% credible interval of the Bayesian model’s prediction. The x-axis represents the latent gametocyte density (log 10 scale), and the y-axis represents the predicted mosquito infection probability.

Asexual parasite density retained a positive residual association with infectivity after conditioning on gametocyte density (β = 0.56, 95% CrI: 0.47–0.64; OR = 1.74). However, pathway decomposition within the joint model indicated that a substantial portion of the parasite–infectivity association operated through gametocyte density. The total association of parasite density with infectivity was strong (TE = 0.94, 95% CrI: 0.85–1.02; OR = 2.55), of which approximately 41% was estimated to operate through the parasite → gametocyte → infectivity pathway (IE = 0.38, 95% CrI: 0.32–0.44; OR = 1.46). Fever was negatively associated with infectivity (β = −0.45, 95% CrI: −0.75 to −0.15; OR = 0.64), indicating reduced transmission potential among symptomatic individuals (Table 2 and Fig 4). Age showed opposing associations with parasite stages. Increasing age was associated with lower latent parasite density (β = −0.03, 95% CrI: −0.04 to −0.02) but higher latent gametocyte density (β = 0.02, 95% CrI: 0.01–0.03). These opposing effects resulted in a small net association between age and infectivity (TE = 0.008, 95% CrI: −0.007 to 0.02). The latent process model also confirmed a strong biological coupling between parasite replication and gametocyte production (γ = 0.45), which showed that each tenfold increase in parasite density was associated with approximately a 2.8-fold increase in gametocyte density. The beta-binomial overdispersion parameter (ϕ = 1.54, 95% CrI: 1.35–1.75) indicated substantial variability in mosquito infection outcomes beyond that expected under a simple binomial model.

### Predicted changes in the proportion of infected mosquitoes

To facilitate interpretation of the joint model, predicted changes in the proportion of infected mosquitoes were calculated under specific contrasts in the model covariates. These contrasts translate the logistic regression results into differences in predicted infection probability while holding other variables constant (detailed description of the result are provided in S1 Text).

Predicted differences associated with a five-year increase in age were generally small across studies (S1 Fig). Across studies, the predicted changes were close to zero, indicating minimal differences in the proportion of infected mosquitoes when age increased by five years. Similarly, predicted changes associated with a tenfold increase in asexual parasite density mediated through gametocyte density were modest across studies (S2 Fig). Although some variation across age groups was observed, predicted differences remained relatively small. Increases in gametocyte density produced substantially larger predicted increases in the proportion of infected mosquitoes (S3 Fig). The magnitude of these differences depended on baseline gametocyte density, with larger increases observed at higher gametocyte levels. These results reinforce the dominant role of gametocyte density in determining mosquito infectivity.

Predicted differences associated with fever status were modest after accounting for parasite and gametocyte densities, consistent with the negative conditional association estimated in the model (S4 Fig). When asexual parasite density was increased while holding gametocyte density constant, predicted differences in infectivity remained modest but were consistently positive across several strata **(S5 Fig)**. This suggests that although much of the association between parasite density and infectivity operates through gametocyte production, asexual parasite density may still capture additional information related to transmission potential that is not fully reflected by measured gametocyte densities. Similarly, residual age effects after conditioning on parasite and gametocyte densities were minimal across studies (S6 Fig).

### Combined effect of parasite and gametocyte density on transmission

Model predictions showed that mosquito infection probability is primarily driven by gametocyte density, but that includes asexual parasite density improves prediction by capturing information about earlier parasite dynamics (preceding gametocyte production) and residual variation (Fig 6). Predicted infectivity increased with gametocyte density across all parasite densities, while higher asexual parasite density further shifted predicted risk upward at a given gametocyte level.

**Fig 6.**
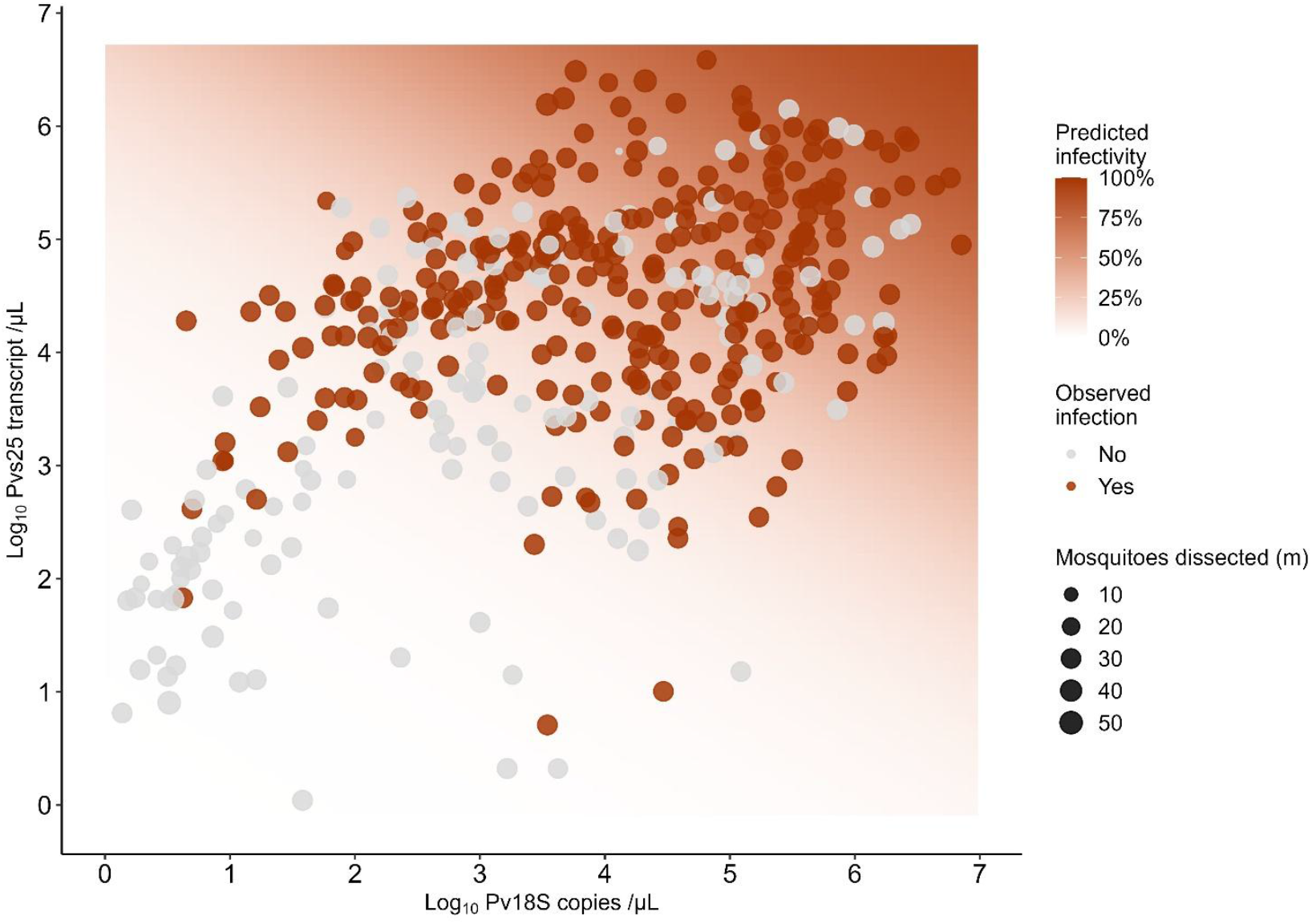
Relationship between parasite density and gametocyte density with observed infections. The y-axis represents PvS25 copies/µL, and the x-axis represents Pv18S copies/µL. The color gradient indicates the likelihood of infection, with red points showing observed infections and grey points showing non-infections. Tile colors indicate predicted mosquito infectivity (white = 0%, dark red = 100%). Points show observed feeding outcomes, colored by infection status and scaled by the number of mosquitoes dissected (m), reflecting the relative precision of each estimate.

Predicted infection probability was highest in the region where both parasite and gametocyte density were high, whereas high parasite density with low gametocyte density was associated with substantially lower predicted infectivity. Observed infections (brown points) clustered in areas of higher predicted infectivity (purple contours), while non-infections (gray points) mainly appeared in low-density areas (yellow contours). This nonlinear interaction shows the interdependence of asexual and sexual stages in driving *P. vivax* transmission.

### Estimates of mean parasite and gametocyte densities

The latent process sub models revealed substantial noise in molecular parasite quantification. The posterior standard deviations for the latent process model (*σ*_*P*,lat_ = 1.44; *σ*_*G*,lat_ = 0.85 on the log10 scale). After accounting for measurement error, the estimated mean latent parasite density (*μ*_*P*_ ) as 3.71 (95% CrI: 3.57–3.85), corresponding to approximately 5,129 parasites/µL (95% CrI: 3,715–7,244) for an individual at the average age of the study participant and the mean latent gametocyte density *(μ*_*G*_ ), defined for an individual with mean-centered age and mean-centered parasite density, was estimated at 4.17 (95% CrI: 4.10–4.25), corresponding to approximately 14,791 Pvs25 transcript /µL (95% CrI: 12,589–17,783) (Table 2).

### Age effects on parasite and gametocyte densities

Age showed opposing associations with asexual parasite and gametocyte densities. The estimated per-year association with latent asexual parasite density was modest (γ_P_age_ = −0.03, 95% CrI: −0.04 to −0.02). When scaled over a 10-year age difference, this corresponds to substantially lower predicted asexual parasite density — for example, a 15-year-old is predicted to have roughly half the asexual parasite density of a 5-year-old (10^−0.03×10^ ≈ 50%).while latent gametocyte density were positively associated with age (γ_G_age_ = 0.02, 95% CrI: 0.01–0.03), corresponding to approximately 60% higher gametocyte transcript levels over the same 10-year age difference ((10^0.02×10^ ≈ 1.58). This opposing age gradient indicates that older individuals may maintain similar or higher transmission marker levels despite lower asexual parasite densities (Table 2 and Fig 3). This represents a conditional association from the joint model and does not necessarily imply higher raw observed gametocyte measurements by age alone.

### Model convergence, performance, and predictive insights

The Bayesian model demonstrated excellent convergence, with Gelman-Rubin diagnostics 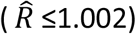 and effective sample sizes (ESS > 3,400) for all parameters, ensuring reliable inference. Posterior predictive checks confirmed that the model accurately reproduced observed mosquito infection distributions. Model adequacy was assessed using posterior predictive checks, while convergence and sampling efficiency were evaluated using trace plots, autocorrelation functions, and effective sample size diagnostics. All diagnostic plots indicated good mixing and convergence of the Markov chains (Supplementary Fig S7–S11). Model performance improved compared to the frequentist approach, as indicated by lower Brier scores (0.073 vs 0.092) and higher AUC (0.83 vs 0.68), demonstrating improved predictive accuracy and uncertainty handling (S4 Table). Compared with the frequentist model considered here, the Bayesian model showed better predictive performance and provided a coherent way to propagate uncertainty across linked biological stages. Predictive accuracy was high (Brier score = 0.073; AUC = 0.83), while the frequentist predictive performance (Brier score = 0.092; AUC = 0.68; S4 Table and S12 Fig). These results suggest that the Bayesian model handled measurement and process uncertainty more effectively in this dataset and may be useful for hypothesis generation.

## Discussion

This study applied a joint Bayesian latent-variable model to quantify relationships among parasite density, gametocyte density, and mosquito infectivity in *Plasmodium vivax* using data from three transmission studies in Ethiopia. By modelling parasite density, gametocyte density, and mosquito infection outcomes within a unified probabilistic framework, the analysis accounted for measurement error, biological dependence between parasite stages, and between-study heterogeneity. The results show that gametocyte density was the strongest predictor of mosquito infectivity, while asexual parasite density retained an additional conditional association with infectivity after accounting for gametocyte density.

The strong association between gametocyte density and mosquito infectivity observed in the joint model (OR=2.3; 95% CrI: 2.2 – 2.5) is consistent existing literature [8, 28]. The estimated relationship between gametocyte density and infection probability showed a sigmoidal pattern, with infection probability increasing rapidly at intermediate gametocyte densities and approaching a plateau at higher densities[29, 30]. This pattern has been reported in studies of malaria transmission that relates gametocyte density to mosquito infection outcomes and reflects the probabilistic nature of transmission events during blood feedings [29, 31]. In this study, predicted infection probability remained low at lower gametocyte densities and increased across an estimated range where infection probability begins to increase around 100 transcripts/µL (95% CrI: 31–292), consistent with experimental observations that transmission probability depends strongly on the density of circulating gametocytes[5, 9, 30].

The joint modeling framework also allowed the parasite–infectivity association to be decomposed into pathway-specific and residual conditional components. The results indicated that a substantial portion of the parasite–infectivity association was explained by the modeled parasite → gametocyte → infectivity pathway, consistent with the biological pathway in which asexual parasite replication precedes gametocyte production[8, 9]. The remaining association between parasite density and infectivity persisted after conditioning on measured gametocyte density, suggesting that parasite density may capture additional information not fully represented by a single gametocyte measurement. Thus, higher asexual parasite density remained associated with higher predicted infectivity even after accounting for measured gametocyte density. Because the data were cross-sectional, measured gametocyte density represents only a single snapshot of a dynamic process. The residual asexual parasite density may be capturing information about recent or impending gametocyte production that is not fully reflected in a single gametocyte measurement, including gametocyte maturity, sex ratio and other temporal dynamics [9, 29]. Similar findings have been reported in studies of *Plasmodium falciparum*, where parasite density influences infectivity primarily through its relationship with gametocyte density[29, 30].

Age showed opposing associations with parasite density and gametocyte density. Increasing age was associated with lower parasite density but slightly higher gametocyte density in the latent process model. These opposing effects resulted in a small overall association between age and mosquito infectivity. Age-related differences in parasite density have been reported in multiple malaria-endemic settings and are commonly attributed to the acquisition of partial immunity that suppresses asexual parasite replication[32, 33]. In contrast, gametocyte carriage has been reported to persist even when parasite densities decline, potentially maintaining transmission potential in older individuals[9, 34]. The joint modeling framework provides a useful way to quantify these opposing associations within a single analytical structure.

In addition to parameter estimates on the log-odds scale, predicted changes in the proportion of infected mosquitoes were calculated under specific contrasts in model covariates. These probability-based summaries provide a biologically interpretable view of model predictions by translating regression parameters into differences in predicted infection probability[35, 36]. Across contrasts, increases in gametocyte density produced the largest predicted differences in the proportion of infected mosquitoes (up to 20%). In some strata, when looking at the residual effects of asexual parasite density, a ten-fold increase in asexual parasite density resulted in a 10% to 15% difference in the proportion infected mosquitos. At high gametocyte and asexual parasite densities, when a fever was present, the expected proportion of infected mosquitos reduced by approximately 10% (in absolute terms). These summaries help translate model coefficients into predicted transmission outcomes under biologically meaningful contrasts [35, 37].

From a methodological perspective, the joint Bayesian latent-variable framework provides a useful approach for analyzing malaria transmission data. First, it allows parasite density, gametocyte density, and mosquito infection outcomes to be modelled jointly rather than as independent processes[35]. Second, the framework explicitly accounts for measurement error in molecular parasite and gametocyte measurements, which can otherwise attenuate estimated associations[35, 38]. Third, the model propagates uncertainty across biological stages, allowing inference about transmission outcomes while accounting for uncertainty in upstream measurements[35, 39]. These features make the approach well suited to infectious disease settings in which multiple interdependent processes are imperfectly observed[39, 40].

Several limitations should be considered when interpreting these findings. The mosquito infectivity data was obtained from membrane feeding assays performed under controlled insectary conditions using *Anopheles arabiensis*. Transmission efficiency and quantitative relationships between parasite densities and mosquito infection outcomes may differ across vector species and ecological settings[41, 42]. In addition, two of the three contributing studies were cross-sectional, which limits inference about temporal processes such as infection duration, relapse dynamics, and longitudinal changes in parasite densities[6, 43]. Although measurement error was explicitly modelled, additional sources of laboratory or assay variability may remain unaccounted for. Finally, unmeasured host or parasite factors, including immunity, parasite genetic diversity, or vector-related variation, may influence mosquito infection outcomes[9].

Despite these limitations, the joint modelling approach provides a framework for linking molecular parasite measurements with transmission outcomes while accounting for uncertainty in multiple biological processes. Although developed here for *P. vivax*, the framework may be applicable to other infectious disease systems where transmission depends on interconnected and imperfectly observed biological stages. By integrating measurement and transmission components within a single probabilistic structure, this approach supports uncertainty-aware inference and clearer interpretation of transmission processes.

## Data availability statement

All data and code used for model fitting and plotting are available on a GitHub repository at https://github.com/AHRI-Malaria/Pvivax-transmission-model.

## Author contributions

**Conceptualization:** Legesse Alamerie Ejigu, John Bradley, Jordache Ramjith,Fitsum G. Tadesse

**Methodology:** Legesse Alamerie Ejigu, John Bradley, Jordache Ramjith

**Software:** Legesse Alamerie Ejigu

**Formal Analyses :** Legesse Alamerie Ejigu, Wakweya Chali, John Bradley, Jordache Ramjith, Chris Drakeley, Teun Bousema, Fitsum G. Tadesse

**Data Curation:** Legesse Alamerie Ejigu, Wakweya Chali

**Visualization:** Legesse Alamerie Ejigu

**Supervision:** John Bradley, Chris Drakeley, Fitsum G. Tadesse, Jordache Ramjith

**Original Draft Preparation:** Legesse Alamerie Ejigu, Fitsum G. Tadesse, John Bradley, Jordache Ramjith, Chris Drakeley

**Writing – Review & Editing:** Legesse Alamerie Ejigu, Wakweya Chali, John Bradley, Jordache Ramjith, Chris Drakeley, Teun Bousema, Fitsum G. Tadesse,

## Supporting information

**S1 Fig**. Predicted differences in the proportion of infected mosquitoes associated with a five-year increase in age.

**S2 Fig**. Predicted differences in the proportion of infected mosquitoes associated with a tenfold increase in parasite density mediated through gametocyte density.

**S3 Fig**. Predicted differences in the proportion of infected mosquitoes associated with a tenfold increase in gametocyte density.

**S4 Fig**. Predicted differences in the proportion of infected mosquitoes associated with fever status.

**S5 Fig**. Predicted differences in the proportion of infected mosquitoes associated with parasite density after conditioning on gametocyte density.

**S6 Fig**. Predicted differences in the proportion of infected mosquitoes associated with age after conditioning on parasite and gametocyte density.

**S7 Fig**. Posterior predictive check for model fit observed vs replicated data.

**S8 Fig**. Trace plots of MCMC samples for model parameters.

**S9 Fig**. Posterior density plots of model parameters.

**S10 Fig**. Autocorrelation diagnostics for MCMC sampling.

**S11 Fig**. Posterior correlation (pairs) plots for infection-related parameters.

**S12 Fig**. Calibration of the prediction for mosquito infection.

**S1 Table**. Summary of Dataset used for model fitting.

**S2 Table**. Prior distributions used in the Bayesian model.

**S3 Table**. Frequentist model results

**S1 Text**. Supplementary methods and results

## References

[1] WHO, “World malaria report 2025, Geneva. World Health Organization. https://www.who.int/teams/global-malaria-programme/reports/world-malaria-report-2024,” 2025.

[2] R. E. Howes et al., “Global Epidemiology of Plasmodium vivax,” The American journal of tropical medicine and hygiene, vol. 95, no. 6 Suppl, pp. 15–34, Dec 28 2016, doi: 10.4269/ajtmh.16-0141.

[3] M. N. Anwar et al., “Mathematical models of Plasmodium vivax transmission: A scoping review,” PLoS Comput Biol, vol. 20, no. 3, p. e1011931, Mar 2024, doi: 10.1371/journal.pcbi.1011931.

[4] F. E. McKenzie et al., “Gametocytemia in Plasmodium vivax and Plasmodium falciparum infections,” J Parasitol, vol. 92, no. 6, pp. 1281–5, Dec 2006, doi: 10.1645/GE-911R.1.

[5] A. Tajebe, G. Magoma, M. Aemero, and F. Kimani, “Detection of mixed infection level of Plasmodium falciparum and Plasmodium vivax by SYBR Green I-based real-time PCR in North Gondar, north-west Ethiopia,” Malar J, vol. 13, p. 411, Oct 18 2014, doi: 10.1186/1475-2875-13-411.

[6] N. J. White, “Determinants of relapse periodicity in Plasmodium vivax malaria,” Malar J, vol. 10, p. 297, Oct 11 2011, doi: 10.1186/1475-2875-10-297.

[7] T. Bousema, L. Okell, I. Felger, and C. Drakeley, “Asymptomatic malaria infections: detectability, transmissibility and public health relevance,” Nat Rev Microbiol, vol. 12, no. 12, pp. 833–40, Dec 2014, doi: 10.1038/nrmicro3364.

[8] J. C. Beier, “Malaria parasite development in mosquitoes,” Annu Rev Entomol, vol. 43, pp. 519–43, 1998, doi: 10.1146/annurev.ento.43.1.519.

[9] T. Bousema and C. Drakeley, “Epidemiology and infectivity of Plasmodium falciparum and Plasmodium vivax gametocytes in relation to malaria control and elimination,” Clin Microbiol Rev, vol. 24, no. 2, pp. 377–410, Apr 2011, doi: 10.1128/CMR.00051-10.

[10] H. Pett et al., “Comparison of molecular quantification of Plasmodium falciparum gametocytes by Pfs25 qRT-PCR and QT-NASBA in relation to mosquito infectivity,” Malar J, vol. 15, no. 1, p. 539, Nov 8 2016, doi: 10.1186/s12936-016-1584-z.

[11] M. F. B. a. S. F. Kitchen, “On the Infectiousness of Patients Infected with Plasmodium Vivax and Plasmodium Falciparum,” The American journal of tropical medicine and hygiene, vol. s1-17, no. 2, 1937, doi: 10.4269/ajtmh.1937.s1-17.253.

[12] J. C. Koella, “On the use of mathematical models of malaria transmission,” Acta Trop, vol. 49, no. 1, pp. 1–25, Apr 1991, doi: 10.1016/0001-706x(91)90026-g.

[13] C. Drakeley, C. Sutherland, J. T. Bousema, R. W. Sauerwein, and G. A. Targett, “The epidemiology of Plasmodium falciparum gametocytes: weapons of mass dispersion,” Trends Parasitol, vol. 22, no. 9, pp. 424–30, Sep 2006, doi: 10.1016/j.pt.2006.07.001.

[14] D. B. Belay et al., “Joint Bayesian modeling of time to malaria and mosquito abundance in Ethiopia,” BMC Infect Dis, vol. 17, no. 1, p. 415, Jun 12 2017, doi: 10.1186/s12879-0172496-4.

[15] Z. G. Dessie, T. Zewotir, H. Mwambi, and D. North, “Modelling of viral load dynamics and CD4 cell count progression in an antiretroviral naive cohort: using a joint linear mixed and multistate Markov model,” BMC Infect Dis, vol. 20, no. 1, p. 246, Mar 26 2020, doi: 10.1186/s12879-020-04972-1.

[16] N. N. McHunu, H. G. Mwambi, T. Reddy, N. Yende-Zuma, and K. Naidoo, “Joint modelling of longitudinal and time-to-event data: an illustration using CD4 count and mortality in a cohort of patients initiated on antiretroviral therapy,” BMC Infect Dis, vol. 20, no. 1, p. 256, Mar 30 2020, doi: 10.1186/s12879-020-04962-3.

[17] E. Hailemeskel et al., “Dynamics of asymptomatic Plasmodium falciparum and Plasmodium vivax infections and their infectiousness to mosquitoes in a low transmission setting of Ethiopia: a longitudinal observational study,” Int J Infect Dis, vol. 143, p. 107010, Jun 2024, doi: 10.1016/j.ijid.2024.107010.

[18] F. G. Tadesse et al., “The Relative Contribution of Symptomatic and Asymptomatic Plasmodium vivax and Plasmodium falciparum Infections to the Infectious Reservoir in a Low-Endemic Setting in Ethiopia,” Clin Infect Dis, vol. 66, no. 12, pp. 1883–1891, Jun 1 2018, doi: 10.1093/cid/cix1123.

[19] T. Eisenberg et al., “Nucleocytosolic depletion of the energy metabolite acetyl-coenzyme a stimulates autophagy and prolongs lifespan,” Cell Metab, vol. 19, no. 3, pp. 431–44, Mar 4 2014, doi: 10.1016/j.cmet.2014.02.010.

[20] A. Gelman, & Hill, J., Data Analysis Using Regression and Multilevel/Hierarchical Models. 2006.

[21] J. K. Kruschke, Doing Bayesian Data Analysis. 2015.

[22] J. T. Lin, D. L. Saunders, and S. R. Meshnick, “The role of submicroscopic parasitemia in malaria transmission: what is the evidence?,” Trends Parasitol, vol. 30, no. 4, pp. 183–90, Apr 2014, doi: 10.1016/j.pt.2014.02.004.

[23] F. G. Tadesse et al., “The shape of the iceberg: quantification of submicroscopic Plasmodium falciparum and Plasmodium vivax parasitaemia and gametocytaemia in five low endemic settings in Ethiopia,” Malar J, vol. 16, no. 1, p. 99, Mar 3 2017, doi: 10.1186/s12936-017-1749-4.

[24] R. McElreath, Statistical Rethinking: A Bayesian Course with Examples in R and Stan (1st ed.). Chapman and Hall/CRC. 2016.

[25] B. Carpenter et al., “Stan: A Probabilistic Programming Language,” J Stat Softw, vol. 76, 2017, doi: 10.18637/jss.v076.i01.

[26] M. Miocevic, O. Gonzalez, M. J. Valente, and D. P. MacKinnon, “A Tutorial in Bayesian Potential Outcomes Mediation Analysis,” Struct Equ Modeling, vol. 25, no. 1, pp. 121–136, 2018, doi: 10.1080/10705511.2017.1342541.

[27] Y. Yuan and D. P. MacKinnon, “Bayesian mediation analysis,” Psychol Methods, vol. 14, no. 4, pp. 301–22, Dec 2009, doi: 10.1037/a0016972.

[28] D. L. Smith and F. E. McKenzie, “Statics and dynamics of malaria infection in Anopheles mosquitoes,” Malar J, vol. 3, p. 13, Jun 4 2004, doi: 10.1186/1475-2875-3-13.

[29] T. S. Churcher et al., “Predicting mosquito infection from Plasmodium falciparum gametocyte density and estimating the reservoir of infection,” Elife, vol. 2, p. e00626, May 21 2013, doi: 10.7554/eLife.00626.

[30] J. Bradley et al., “Predicting the likelihood and intensity of mosquito infection from sex specific Plasmodium falciparum gametocyte density,” Elife, vol. 7, May 31 2018, doi: 10.7554/eLife.34463.

[31] P. Schneider et al., “Submicroscopic Plasmodium falciparum gametocyte densities frequently result in mosquito infection,” The American journal of tropical medicine and hygiene, vol. 76, no. 3, pp. 470–4, Mar 2007. [Online]. Available: https://www.ncbi.nlm.nih.gov/pubmed/17360869.

[32] J. K. Baird, “Evidence and implications of mortality associated with acute Plasmodium vivax malaria,” Clin Microbiol Rev, vol. 26, no. 1, pp. 36–57, Jan 2013, doi: 10.1128/CMR.00074-12.

[33] I. Rodriguez-Barraquer et al., “Quantification of anti-parasite and anti-disease immunity to malaria as a function of age and exposure,” Elife, vol. 7, Jul 25 2018, doi: 10.7554/eLife.35832.

[34] C. Koepfli and G. Yan, “Plasmodium Gametocytes in Field Studies: Do We Measure Commitment to Transmission or Detectability?,” Trends Parasitol, vol. 34, no. 5, pp. 378-387, May 2018, doi: 10.1016/j.pt.2018.02.009.

[35] J. B. C. Andrew Gelman, Hal S. Stern, David B. Dunson, Aki Vehtari, Donald B. Rubin, Data Analysis Using Regression and Multilevel/Hierarchical Models. 2006.

[36] D.B. Norton EC, Maciejewski ML., “Marginal Effects—Quantifying the Effect of Changes in Risk Factors in Logistic Regression Models,” JAMA, 2019, doi: doi:10.1001/jama.2019.1954.

[37] S. Greenland, Schlesselman, J.J. and Criqui, M.H., “The Fallacy of Employing Standardized Regression Coefficients and Correlations as Measures of Effect.,” American Journal of Epidemiology, vol. 125, no. 2, pp. 349–350, 1986, doi: 10.1093/oxfordjournals.aje.a114539.

[38] R. J. Carroll, Ruppert, D., Stefanski, L.A., & Crainiceanu, C.M, Measurement Error in Nonlinear Models: A Modern Perspective, Second Edition (2nd ed.). 2006.

[39] R. McElreath, Statistical Rethinking: A Bayesian Course with Examples in R and STAN (2nd ed.). 2020.

[40] H. Heesterbeek et al., “Modeling infectious disease dynamics in the complex landscape of global health,” Science, vol. 347, no. 6227, p. aaa4339, Mar 13 2015, doi: 10.1126/science.aaa4339.

[41] T. Bousema and C. Drakeley, “Determinants of Malaria Transmission at the Population Level,” Cold Spring Harb Perspect Med, vol. 7, no. 12, Dec 1 2017, doi: 10.1101/cshperspect.a025510.

[42] A. Ahmad et al., “Infectivity of patent Plasmodium falciparum gametocyte carriers to mosquitoes: establishing capacity to investigate the infectious reservoir of malaria in a low-transmission setting in The Gambia,” Trans R Soc Trop Med Hyg, vol. 115, no. 12, pp. 1462–1467, Dec 2 2021, doi: 10.1093/trstmh/trab087.

[43] K. E. Battle et al., “Mapping the global endemicity and clinical burden of Plasmodium vivax, 2000-17: a spatial and temporal modelling study,” Lancet, vol. 394, no. 10195, pp. 332–343, Jul 27 2019, doi: 10.1016/S0140-6736(19)31096-7.

